# Inhibition of ApoE4 Endocytosis with LDLR-LA Peptides

**DOI:** 10.64898/2026.05.20.726622

**Authors:** Sang Ho Park, Gregory-Neal Gomes, Brittney A. Beyer, Zachary A. Levine

## Abstract

The Apolipoprotein E4 (ApoE4) genotype is the most significant genetic risk factor for late-onset Alzheimer’s disease (AD). A key driver of ApoE4 cellular toxicity is the endo-lysosomal burden resulting from the excessive receptor-mediated uptake of ApoE4 lipoparticles. The high-affinity interaction between lipidated ApoE4 and the Low-Density Lipoprotein Receptor (LDLR) saturates the cellular degradation machinery, correlating with lysosomal alkalinization, lipid accumulation, and cell death. To target this critical interaction interface, which consists of 7 tandem ligand-binding type-A (LA) modules in the human LDLR, we present the design and evaluation of recombinant LDLR minireceptors comprising combinations of these LA modules to competitively antagonize ApoE4 endocytosis. We observe a distinct isoform-dependent uptake dynamic across multiple central nervous system (CNS) cell models, with ApoE4 showing significantly greater total intracellular accumulation than ApoE2. Furthermore, engineered LA peptides selectively bind ApoE4 over human serum LDL and differentially inhibit its uptake, revealing a distinct structural efficacy hierarchy of LA3456 ∼ LA345 > LA456 ∼ LA45 >> LA34. We establish the resilience of the LA45 minireceptor under physiological serum conditions and identify LA345 as the most stable truncated construct *in vitro*. Notably, molecular tagging orientation is critical for therapeutic engineering; C-terminal tagging completely preserves the inhibitory function of the minireceptors, whereas N-terminal tagging drastically reduces it. These findings provide a framework for scalable, deliverable inhibition of the ApoE4-LDLR interaction as a potential therapeutic target to mitigate endo-lysosomal accumulation in AD.

## 1. Introduction

The increasing global prevalence of Alzheimer’s disease (AD) highlights an urgent need for disease-modifying therapies. While the amyloid cascade hypothesis has guided therapeutic development for decades, the major molecular risk factor for late-onset AD, the expression of the ε4 allele of the apolipoprotein E (*APOE*) gene, remains a largely un-drugged target (Martens et al., 2022; Windham & Cohen, 2023).

ApoE is a 34-kDa glycoprotein that functions as the brain’s primary lipid and cholesterol transporter. It plays an essential role in redistributing lipids among CNS cells to maintain membrane homeostasis, facilitate synapto-dendritic connection repair, and clear toxic debris (Cantuti-Castelvetri et al., 2018; Bales et al., 1997). The *APOE* gene encodes three major human isoforms: ApoE2, ApoE3, and ApoE4. These isoforms differ by single amino acid substitutions at positions 112 and 158. While the ApoE2 allele is generally neuroprotective, the ApoE4 allele is present in a vast majority of late-onset AD patients and significantly accelerates the age of disease onset (Shinohara et al., 2020; Li et al., 2020).

Clinical evidence demonstrates that carriers of the ApoE4 allele face a drastically increased risk of developing late-onset AD, heavily exacerbating tau-mediated neurodegeneration and disrupting homeostatic functions in glia (Shi et al., 2017; Fernandez et al., 2019; Ferrari-Souza et al., 2023). This structural variance significantly affects the binding affinity of ApoE for cell surface receptors (Chen et al., 2011; Chen et al., 2021). The Low-Density Lipoprotein Receptor (LDLR) family exhibits the highest affinity for lipidated ApoE among brain receptors, and this interaction is highly isoform-dependent (Brown & Goldstein, 1983; Lane-Donovan & Herz, 2017). Specifically, ApoE4 exhibits an exceptionally strong binding affinity for LDLR compared to the weaker affinities of ApoE3 and ApoE2 (Weisgraber et al., 1982; Yamamoto et al., 2008; Guo et al., 2025). Overexpression of LDLR has been shown to drastically alter the cellular uptake of ApoE, reducing overall extracellular levels and modulating Aβ clearance (Kim et al., 2009; Basak et al., 2012).

Because of this high affinity, ApoE4 lipoparticles are rapidly and excessively endocytosed by glia and neurons, driving a massive influx of heavy lipid cargoes into the cell. Once endocytosed, these ApoE lipoparticles traffic directly to the endo-lysosomal system. The accumulation of these molecules saturates the lysosomes, leading to lysosomal dysfunction, lipid accumulation, and downstream cellular toxicity (Guo et al., 2025). This phenomenon correlates with the intracellular accumulation of lipid droplets, unesterified cholesteryl esters, lipofuscin, and aggregated tau (Haney et al., 2024; Feringa & van der Kant, 2021; Seehafer & Pearce, 2006). Given that this mechanism of intracellular toxicity is fundamentally driven by the initial receptor binding and internalization step, this interaction acts as a major biochemical bottleneck.

The native human LDLR utilizes specific clusters of cysteine-rich complement-type ligand-binding (LA) repeats to recognize ApoE (Beglova & Blacklow, 2005). Previous biophysical studies have mapped the interaction interface, demonstrating that minimal multi-domain units, specifically LA4 and LA5, are critical for recognizing ApoE in its physiologically relevant lipid-bound form (Fisher et al., 2004; Guttman et al., 2010a; Guttman et al., 2010b; Guttman & Komives, 2011). Based on these structural principles, this study aims to engineer and validate recombinant LDLR minireceptors as competitive inhibitors. By utilizing varying constructs of LDLR-LA modules, we hypothesized that we could selectively block ApoE4 lipoparticles at the cell surface. Preventing this excessive endocytosis rescues the endo-lysosomal system from becoming overburdened (Guo et al., 2025), highlighting targeted inhibition of this pathway as a potential therapeutic strategy for ApoE-related neurodegenerative diseases.

## 2. Materials and methods

### 2.1 Design, expression, and purification of LDLR-LA minireceptors

Recombinant Low-Density Lipoprotein Receptor (LDLR) minireceptors were rationally designed based on the native human LDLR ectodomain sequence (UniProtKB: P01130). The native receptor utilizes seven tandem ligand-binding type-A (LA) modules at its N-terminus. Each LA domain consists of approximately 40 amino acids stabilized by three disulfide bonds. For structural screening, specific truncated combinations of these repeats—encompassing the third, fourth, fifth, and sixth modules—were synthesized. The resulting engineered variants included LA34, LA45, LA345, LA456, and LA3456. The corresponding amino acid sequences of these peptides, relative to the mature form of human LDLR, are K86-G171, V124-A211, K86-A211, V124-L254, and K86-L254, respectively.

To ensure solubility and facilitate subsequent affinity purification, the LA peptides were generated as expression constructs featuring an N-terminal His6-SUMO tag. The constructs utilized a pET-31b(+) vector and were transformed into SHuffle T7 competent *E. coli* (New England Biolabs) to facilitate disulfide bond formation in the cytoplasm. Following bacterial expression at 30°C, the His6-SUMO-LA peptides were isolated using Immobilized Metal Affinity Chromatography (IMAC). The His6-SUMO tag was cleaved using His-tagged SUMO protease (Trialtus Bioscience) at a protein-to-protease ratio of 100:1 (w/w) at 4°C for 16–20 hours within a 3-kDa dialysis membrane against refolding buffer (20 mM HEPES, 150 mM NaCl, 5 mM cysteine, 1 mM cystine, 5 mM CaCl_2_, pH 7.4). The cleaved peptides were separated from the His6-SUMO tag using reverse IMAC purification. The IMAC flowthrough containing the LA peptides was further purified by fast protein liquid chromatography (FPLC) using a HiLoad 26/600 Superdex 200 size-exclusion chromatography (SEC) column (Cytiva) equilibrated in SEC buffer (20 mM HEPES, 150 mM NaCl, 1 mM CaCl_2_). The purity and expected molecular weights of the isolated LA peptide combinations were verified via SDS-PAGE and LC/MS-TOF, confirming highly pure monomeric fractions ranging between 10 and 25 kDa depending on the domain length.

To systematically investigate the steric impact of molecular tags on the minireceptors’ ApoE4-binding capacity, specific modified variants of the LA45 minireceptor were additionally synthesized. These included two N-terminal extended constructs (His6-SUMO-linker-LA45 and His6-linker-LA45) and two C-terminal tagged constructs utilizing standard epitope tags (LA45-HA and LA45-V5). A bait LA peptide, LA3456-linker-His6, was also generated using the same procedure. The amino acid sequence of the flexible linker is GSGGSG.

The structural stability of the purified LA peptides (LA34, LA45, LA345, LA456, and LA3456) were systematically evaluated. Aliquots of the peptides were stored in SEC buffer at 4°C for two weeks. The stored samples were then analyzed alongside freshly prepared control batches via SDS-PAGE. To thoroughly assess disulfide bond integrity and proper folding maintenance, the gels were run under both reduced and unreduced denaturing conditions.

### 2.2 Preparation and validation of fluorescent ApoE lipoparticles

Recombinant human ApoE isoforms (ApoE2, ApoE3, and ApoE4) were individually expressed and purified using a standard protocol (Newhouse & Weisgraber, 2011). Purified ApoE isoforms were lipidated using 1-palmitoyl-2-oleoyl-glycero-3-phosphocholine (POPC) and 1,2-dioleoyl-sn-glycero-3-phosphoethanolamine-N-TopFluor™ AF594 (PE-AF594) to form biologically relevant, stable lipoparticles. Both lipids were obtained from Avanti Research. For lipidation, approximately 10 uM ApoE was added to POPC and PE-AF594 solubilized in 100 mM sodium cholate at an ApoE:POPC:PE-AF594 molar ratio of 1:85:5 in PBS. The mixture was incubated for 2 hours at 4°C in the dark. Subsequently, the solution was dialyzed within a 10-kDa dialysis membrane at 4°C for 3 days in the dark against PBS, with buffer changes performed twice daily.

Particle uniformity and the successful integration of the ApoE protein with the AF594-labeled lipids were validated via FPLC/SEC. Elution profiles were generated by continuously mapping the retention volume (mL) against dual optical absorbance at 280 nm (confirming protein presence) and 590 nm (confirming the AF594 fluorophore). Robust, synchronized co-elution peaks validated the structural integrity of the ApoE2, ApoE3, and ApoE4 lipoparticles prior to cellular application.

### 2.3 IMAC pull-down assays

To biochemically confirm the direct physical interaction and selectivity between the engineered minireceptors and ApoE lipoparticles, IMAC pull-down assays were performed. The His-tagged LA3456 construct (LA3456-H6) was utilized as the stationary bait phase.

To assess targeted binding, LA3456-H6 was co-incubated with purified ApoE4 lipoparticles. To assess selectivity and cross-reactivity, LA3456-H6 was separately co-incubated with native human serum LDL (Invitrogen). Following co-incubation, the complexes were passed through the IMAC resin. Non-bound proteins were collected in the flowthrough and wash fractions. The bound complexes were subsequently eluted using a highly specific, progressive buffer gradient of 2 mM, 4 mM, 6 mM, 8 mM, and 10 mM TCEP/EDTA. All resulting fractions were analyzed via SDS-PAGE to track the co-elution profiles of ApoE4 (∼37 kDa) against potential competitive targets, such as ApoB and Human Serum Albumin (HSA).

### 2.4 Cell culture models and live-cell endocytosis assays

*In vitro* endocytosis and inhibition evaluations were conducted using three distinct central nervous system (CNS) cell lines: wild-type primary human astrocytes (ScienCell #1800), ApoE knockout (KO) iAstrocytes (iAstrocytes-KO) generated from APOE-KO iPSCs (AlStem; StemCell Technologies StemDiff Kits), and human microglial clone 3 (HMC3) cells (ATCC CRL-3304). Cells were maintained in standard appropriate culture media under standard incubator conditions (37°C, 5% CO_2_). For the live-cell assays, primary astrocytes and HMC3 cells were seeded at a density of 10,000 cells per well onto 96-well optical plates (Revvity 6055302), while iAstrocytes-KO were seeded at a density of 5,000 cells per well. All cells were allowed to adhere overnight.

The following day, the culture medium was replaced with serum-free medium. To establish baseline uptake metrics, cells were incubated with 100 nM of the AF594-labeled ApoE2, ApoE3, or ApoE4 lipoparticles. For competitive inhibition assays, exogenous LDLR-LA minireceptors were introduced into the culture media concurrently with the ApoE lipoparticles across a broad dose-escalation gradient (0, 0.1, 0.2, 0.4, 0.8, 1.6, and 3.2 µM). To rigorously assess the minireceptors’ efficacy under physiological constraints and specifically evaluate the effect of serum on LA45 inhibition efficacy, parallel assays were conducted comparing standard serum-free conditions (-FBS) against media supplemented with 10% fetal bovine serum (+FBS).

Endocytosis was tracked via live-cell fluorescence microscopy (Incucyte) over a continuous 24-hour time course, with specific snapshot imaging and analysis occurring at 4-hour intervals. Following the initial 24-hour endocytosis assay, the medium was replaced with serum-containing medium, and the intracellular localization of the ApoE lipoparticles was monitored over the subsequent three days. Endocytic volume was quantified by measuring the total red object integrated fluorescence intensity (FI) per field of view using automated image analysis software.

## 3. Results

### 3.1 Validation of recombinant LA minireceptors and ApoE lipoparticles

To effectively intercept ApoE4 particles in the extracellular space, we sought to mimic the binding capability of the native LDLR by generating truncated variants of its LA modules (Figure 1A). The His6-SUMO tagged LA constructs were robustly expressed in the soluble fraction in Shuffle T7 *E coli*. Subsequent purification using IMAC and FPLC/SEC yielded high-purity isolates of the varied LA peptide combinations (LA34, LA45, LA345, LA456, and LA3456) (Figure 1B). Upon SDS-PAGE, the reduced peptides migrated faster than the non-reduced peptides, demonstrating that the intra-module disulfide bonds remained intact under non-reducing conditions (Figure S1). The correct folding of these peptides, characterized by three disulfide bonds per module, was also validated by LC/MS-TOF.

**Figure 1.**
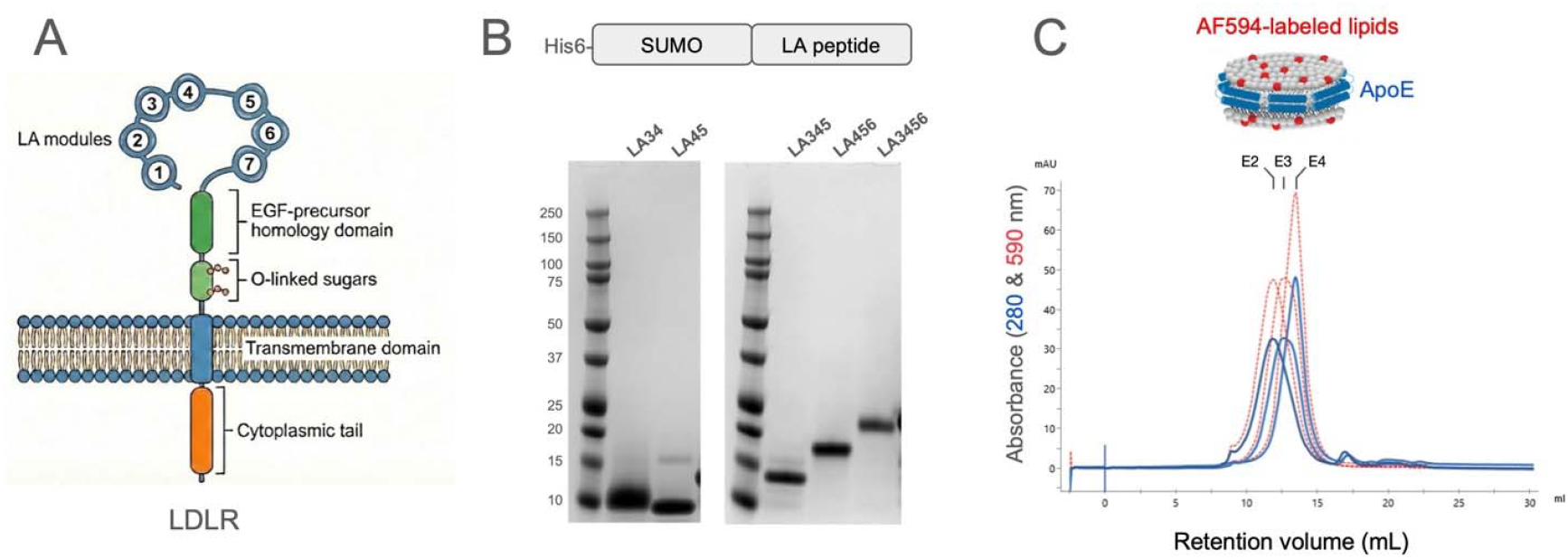
Generation of LDLR-LA peptides and ApoE lipoparticles. (A) Schematic drawing of the human Low-Density Lipoprotein Receptor (LDLR). The extracellular N-terminus contains seven ligand-binding type-A (LA) modules. (B) Design and purification of recombinant LA peptides: The upper schematic illustrates the engineered expression construct, which features an N-terminal His6-SUMO tag fused to the target LA peptide. The lower panels display SDS-PAGE confirming high-purity isolation of the varied truncated LA peptide combinations: LA34, LA45, LA345, LA456, and LA3456. (C) Lipoparticle synthesis and validation: The top drawing depicts the assembly of ApoE with AF594-labeled lipids to form trackable, fluorescent ApoE lipoparticles. The bottom graph shows the size exclusion chromatography (SEC) elution profiles for the synthesized lipoparticles across different isoforms (E2, E3, and E4). The profile maps retention volume (mL) against dual absorbance (280 and 590 nm) to validate successful lipidation and proper particle assembly.

ApoE lipoparticles were assembled with AF594-labeled lipids to form trackable, fluorescent complexes (Figure 1C). Size exclusion chromatography validated the successful lipidation, demonstrating robust co-elution peaks matching absorbance at 280 nm and 590 nm across all three ApoE isoforms (E2, E3, and E4). Notably, the SEC elution profile indicate that the size of ApoE lipoparticles are isoform-dependent, with ApoE2 lipoparticles being the largest and ApoE4 lipoparticles the smallest. This size difference is consistent with observations reported by other groups (Hubin et al., 2019, Ralhan et al., 2026). Furthermore, no significant differences in size and behavior were observed between lipoparticles containing ApoE purified from *E coli* versus those utilizing ApoE derived from mammalian cells.

Manufacturing viable peptide therapeutics requires robust shelf stability. SDS-PAGE analysis of peptides stored at 4°C for over two weeks revealed distinct degradation profiles. Among the synthesized variants, LA345 demonstrated the highest stability, displaying intact structural bands under both reduced and unreduced conditions, whereas LA456 showed signs of oligomerization under unreduced conditions (Figure S3).

### 3.2 Selective binding capacity of recombinant LA minireceptors

Figure 2A illustrates the co-elution of the C-terminally His6-tagged LA3456 (LA3456-H6) construct and ApoE4 lipoparticles from the IMAC resin, demonstrating their direct interaction in solution. It is crucial that a viable therapeutic decoy receptor must exhibit specificity to avoid disrupting essential peripheral lipid transport. To determine whether the minireceptors exhibit selective binding for ApoE4 lipoparticles, we assessed the potential cross-reactivity of the LA3456-H6 construct by co-incubating it with human serum LDL, thereby evaluating its capacity to discriminate between ApoE and ApoB (Figure 2B). The primary structural components of LDL and serum, ApoB (>250 kDa) and human serum albumin (HSA, ∼60–70 kDa), were observed in the flowthrough and wash fractions and failed to co-elute with the LA3456 minireceptor. This confirms the targeted selectivity of LA3456 for ApoE4 over standard circulating LDL.

**Figure 2.**
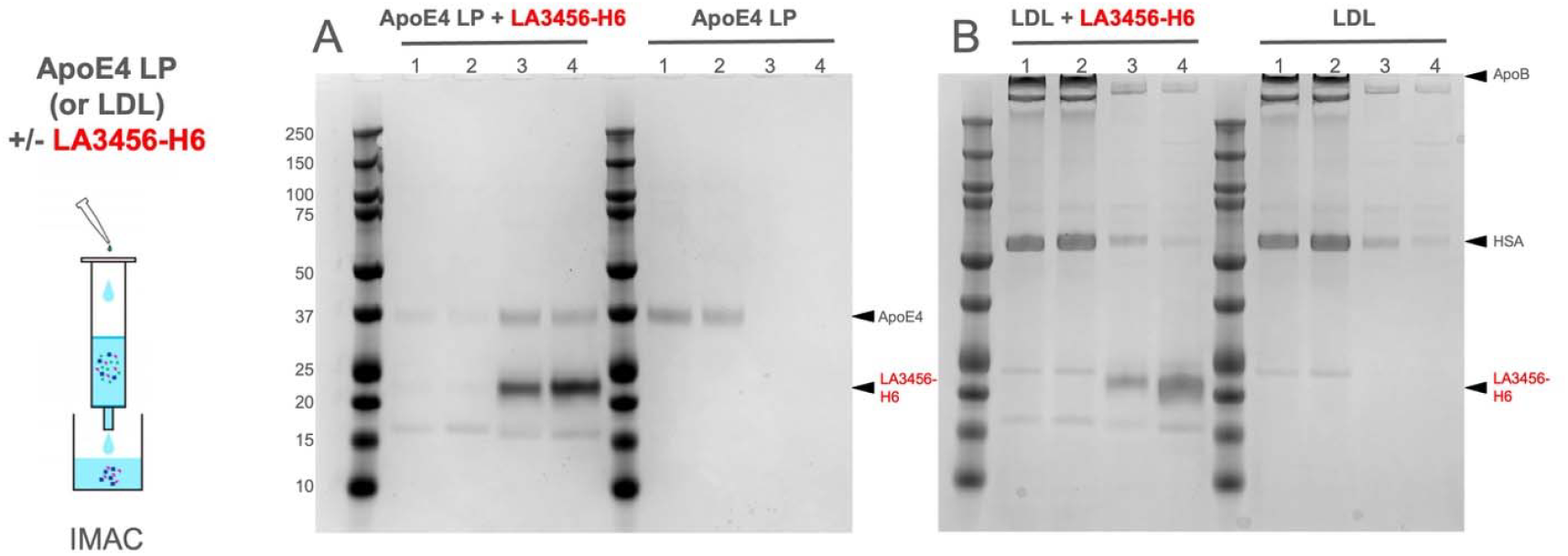
LA3456 selectively binds ApoE4 lipoparticles over human serum LDL. Immobilized Metal Affinity Chromatography (IMAC) pull-down assays were performed to evaluate the binding specificity of the His-tagged LA3456 minireceptor. For all gels, the lanes correspond to the following fractions: lane 1, flowthrough; lane 2, wash; lanes 3 and 4, elution. (A) Interaction with ApoE4 lipoparticles: The left gel demonstrates the co-incubation of ApoE4 lipoparticles (ApoE4 LP) with LA3456-H6. Both ApoE4 (∼37 kDa) and LA3456-H6 (∼20 kDa) successfully co-elute in lanes 3 and 4, indicating a strong binding interaction. The right gel serves as a negative control with ApoE4 LP alone, confirming that ApoE4 does not non-specifically bind to the IMAC column. (B) Interaction with Human Serum LDL: The left gel evaluates the cross-reactivity of LA3456-H6 when co-incubated with human serum LDL. The primary structural components of LDL and serum, ApoB (>250 kDa) and Human Serum Albumin (HSA, ∼60-70 kDa), appear strictly in the flowthrough and wash fractions (lanes 1 and 2) and do not co-elute with LA3456-H6 in lanes 3 and 4. The right gel is a control containing only LDL. This confirms the targeted selectivity of the LA3456 minireceptor for ApoE4 over standard circulating LDL.

Having established this targeted selectivity, we then performed IMAC pull-down assays to confirm the robust physical interaction and optimize the separation of ApoE lipoparticles from the bait construct. Co-incubation of ApoE4 lipoparticles with the LA3456-H6 construct, followed by a progressive buffer gradient elution utilizing 2 to 10 mM TCEP/EDTA, allowed for the optimization of the pull-down of ApoE lipoparticles (Figure 3). ApoE4 lipoparticles were efficiently dissociated from the bait construct and eluted in the presence of 4–6 mM TCEP/EDTA, whereas the LA3456-H6 bait remained bound to the resin and only eluted upon the addition of 300 mM imidazole. Furthermore, fluorescently labeled AF594 lipids were successfully detected at the bottom of the SDS-PAGE gel within the ApoE4 elution fractions, demonstrating the recovery of intact ApoE4 lipoparticles rather than lipid-free ApoE protein. This method facilitates the efficient purification of functional ApoE4 lipoparticles under mild conditions without inducing denaturation, which is critical for preserving their structural integrity for sensitive downstream experiments.

**Figure 3.**
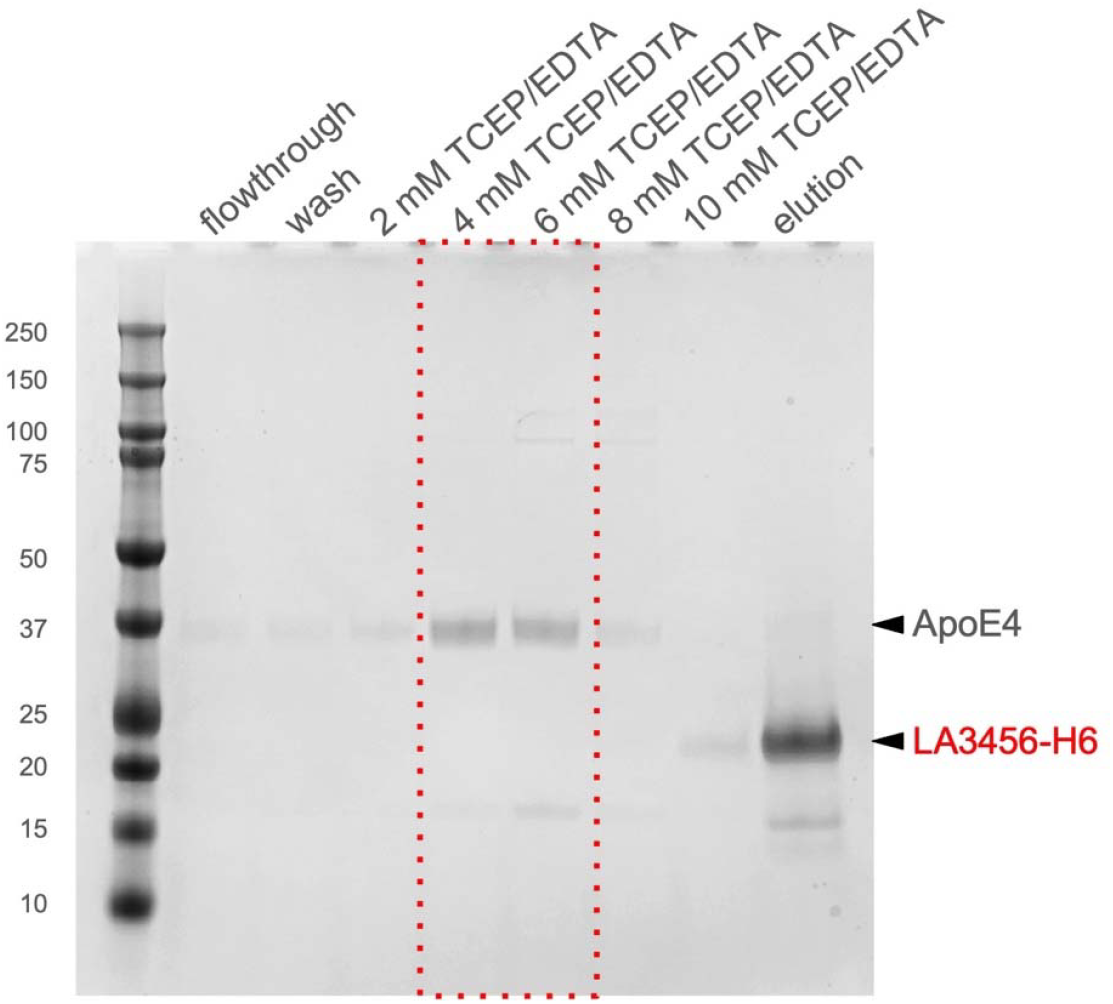
LA3456 successfully pulls down ApoE4 lipoparticles. IMAC was utilized to assess the physical interaction between ApoE4 lipoparticles (ApoE4 LP) and the His-tagged LA3456 minireceptor (LA3456-H6). The SDS-PAGE gel tracks the protein fractions throughout the purification process. Elution of the bound complexes was achieved using a progressive buffer gradient of 2 mM, 4 mM, 6 mM, 8 mM, and 10 mM TCEP/EDTA. The final elution fraction confirms the co-elution of ApoE4 (indicated near the 37 kDa marker) and LA3456-H6 (indicated near the 20 kDa marker), demonstrating a successful binding interaction between the minireceptor and the lipoparticle.

### 3.3 Isoform-dependent endocytosis in CNS cells

To understand how the different variants are internalized, we established the baseline endocytic profiles of fluorescently labeled ApoE2, ApoE3, and ApoE4 lipoparticles across three distinct CNS cell models: wild-type primary human astrocytes, human microglial clone 3 (HMC3) cells, and ApoE knockout astrocytes (iAstrocytes-KO). When these cells were exposed to 100 nM of the respective lipoparticles over a continuous 24-hour time course, a clear isoform dependency was observed that was highly consistent across all evaluated cell types.

Live-cell imaging demonstrated a progressive and massive accumulation of ApoE3 and ApoE4 lipoparticles within the intracellular space. As detailed in the supplemental high-resolution time-course tracking (Figure S1), early endocytic events for ApoE3 and ApoE4 were visible by 4 hours and evolved into robust intracellular accumulation by 12 and 24 hours (Figure 4A–I). In contrast, ApoE2 lipoparticles exhibited visually minimal cellular uptake throughout the entire duration of the assay across all replicates.

**Figure 4.**
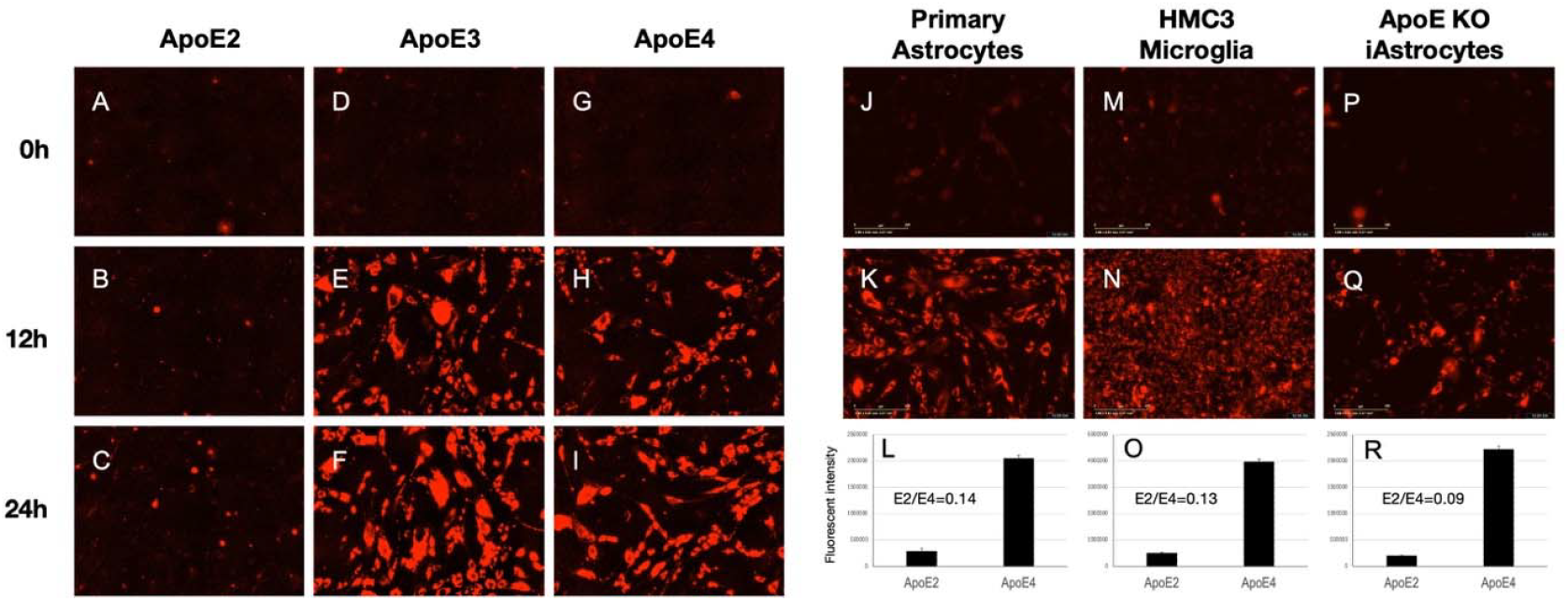
Isoform-dependent endocytosis of ApoE lipoparticles in various CNS cell types. Representative fluorescence microscopy images tracking the internalization of AF594-labeled ApoE lipoparticles across three distinct CNS cell models: Primary Astrocytes, HMC3 Microglia, and ApoE KO iAstrocytes. (A-J) Isoform comparison: Cells were incubated with either ApoE2, ApoE3, or ApoE4 lipoparticles. Cellular uptake was visually monitored across a time gradient at 0h, 12h, and 24h post-treatment. The image panels demonstrate a progressive, time-dependent accumulation of ApoE3 and ApoE4, whereas ApoE2 uptake remains visually minimal throughout the 24-hour observation period. (J-R). Quantification of endocytosis: The bar graphs quantify the total fluorescent intensity for ApoE2 versus ApoE4 at the 24-hour endpoint. The severe isoform bias favoring ApoE4 entry is emphasized by the low calculated E2/E4 uptake ratios across all cell lines: E2/E4=0.14 for Primary Astrocytes, E2/E4=0.13 for HMC3 Microglia, and E2/E4=0.09 for ApoE KO iAstrocytes.

Quantitative analysis of the total integrated fluorescence intensity at the 24-hour endpoint emphasized a severe isoform bias favoring ApoE4 entry (Figure 4J–R). The overall endocytic volume strictly followed a hierarchical rank order of ApoE4 ∼ ApoE3 >> ApoE2. This significant differential was quantitatively reflected by extremely low E2/E4 uptake ratios across the diverse cell lines—specifically 0.14 in primary astrocytes, 0.13 in HMC3 microglia, and 0.09 in iAstrocytes-KO (Figure 4L, 4O, 4R). These findings underscore a pronounced hyper-accumulation of the pathogenic ApoE4 isoform compared to its protective ApoE2 counterpart within these CNS cell models.

### 3.4 Endocytosis Inhibition hierarchy of LA minireceptors

We subsequently tested the ability of the varied LA peptide length combinations to actively block ApoE4 endocytosis in living primary astrocytes (Figure 5). Using a dose-escalation matrix ranging from 0 to 3.2 µM, we tracked the reduction of intracellular red fluorescence to determine the precise inhibitory efficacy of each construct. As visualized in the fluorescence imaging matrix (Figure 5A), the administration of these minireceptors across the concentration gradient resulted in a clear, dose-dependent attenuation of the ApoE4 lipoparticle signal for the majority of the variants tested.

**Figure 5.**
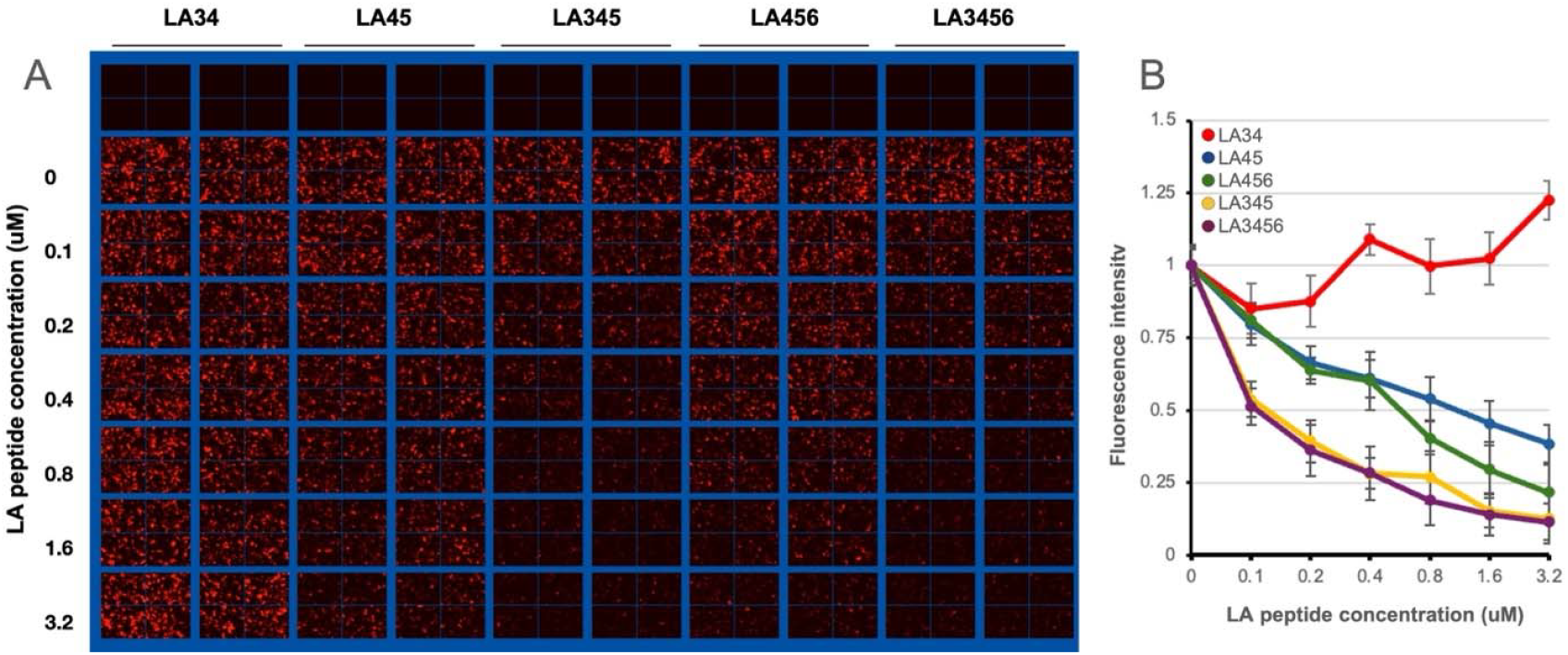
LA peptides differentially inhibit ApoE4 lipoparticle endocytosis. (A) Dose-dependent fluorescence imaging: Representative fluorescence microscopy grid depicting the inhibition of AF594-labeled ApoE4 lipoparticle uptake. Cells were incubated with ApoE4 lipoparticles in the presence of varying concentrations (0, 0.1, 0.2, 0.4, 0.8, 1.6, and 3.2 µM) of the engineered LA minireceptors: LA34, LA45, LA345, LA456, and LA3456. A visual dose-dependent reduction in intracellular red fluorescence is evident for active constructs as peptide concentration increases. (B) Quantification of inhibition hierarchy: Quantitative dose-response curves plotting the normalized cellular fluorescence intensity against LA peptide concentration. The data establishes a distinct inhibitory hierarchy among the modules: the multi-domain constructs LA3456 (purple) and LA345 (yellow) demonstrate the highest potency, rapidly suppressing uptake at low concentrations. LA456 (green) and LA45 (blue) show moderate, steady inhibition across the concentration gradient. Conversely, the truncated LA34 peptide (red) exhibits no significant inhibitory effect even at the maximum dose of 3.2 µM, highlighting the structural necessity of the complete LA4-LA5 tandem for effective ApoE4 antagonism.

The peptides displayed highly differential inhibitory capabilities, establishing a clear endocytosis inhibition hierarchy: LA3456 _≈_ LA345 > LA456 _≈_ LA45 >> LA34 (Figure 5B). The multi-domain LA3456 and LA345 constructs demonstrated the highest potency, rapidly suppressing ApoE4 uptake at relatively low concentrations. Specifically, these longer constructs achieved a half-maximal inhibitory concentration (IC_50_) estimated near 0.1 µM. The LA456 and LA45 constructs showed moderate, steady inhibition across the concentration gradient, reaching their half-maximal inhibition between 0.4 and 0.8 µM. Crucially, the two-domain LA34 peptide was uniquely ineffective, exhibiting no significant inhibitory effect even at the highest dose of 3.2 µM. This functional contrast highlights that the complete LA4-LA5 tandem is structurally indispensable for effective ApoE4 antagonism.

To evaluate the potential effect of the specific fluorescent dye on ApoE lipoparticle behavior, we also tested the endocytosis of ApoE4 lipoparticles containing AF488-labeled lipids (green fluorescence) in human astrocytes in the absence and presence of the minireceptors. We observed the same robust uptake profiles of the AF488-ApoE4 lipoparticles in the untreated control group, with progressive intracellular accumulation clearly visible from 10 minutes through 18 hours of continuous live-cell monitoring (Figure S2, top row). Consistent with our quantitative data, co-incubation with LA34 failed to inhibit this endocytosis, mirroring the heavy uptake profile of the untreated control (Figure S2, middle row). By contrast, the introduction of LA45 effectively intercepted the lipoparticles, resulting in a dramatic and sustained inhibition of their cellular uptake across the entire observed time course (Figure S2, bottom row). This orthogonal validation confirms that the observed uptake profiles and the subsequent effective inhibition by LA45 are driven by specific receptor-ligand dynamics rather than dye-related artifacts.

### 3.5 Inhibitory efficacy under physiological serum conditions

Because therapeutic applications require efficacy in complex physiological environments rich in competing lipids and serum proteins, we evaluated the performance of the LA45 peptide in the presence of 10% fetal bovine serum (FBS) (Figure 6).

**Figure 6.**
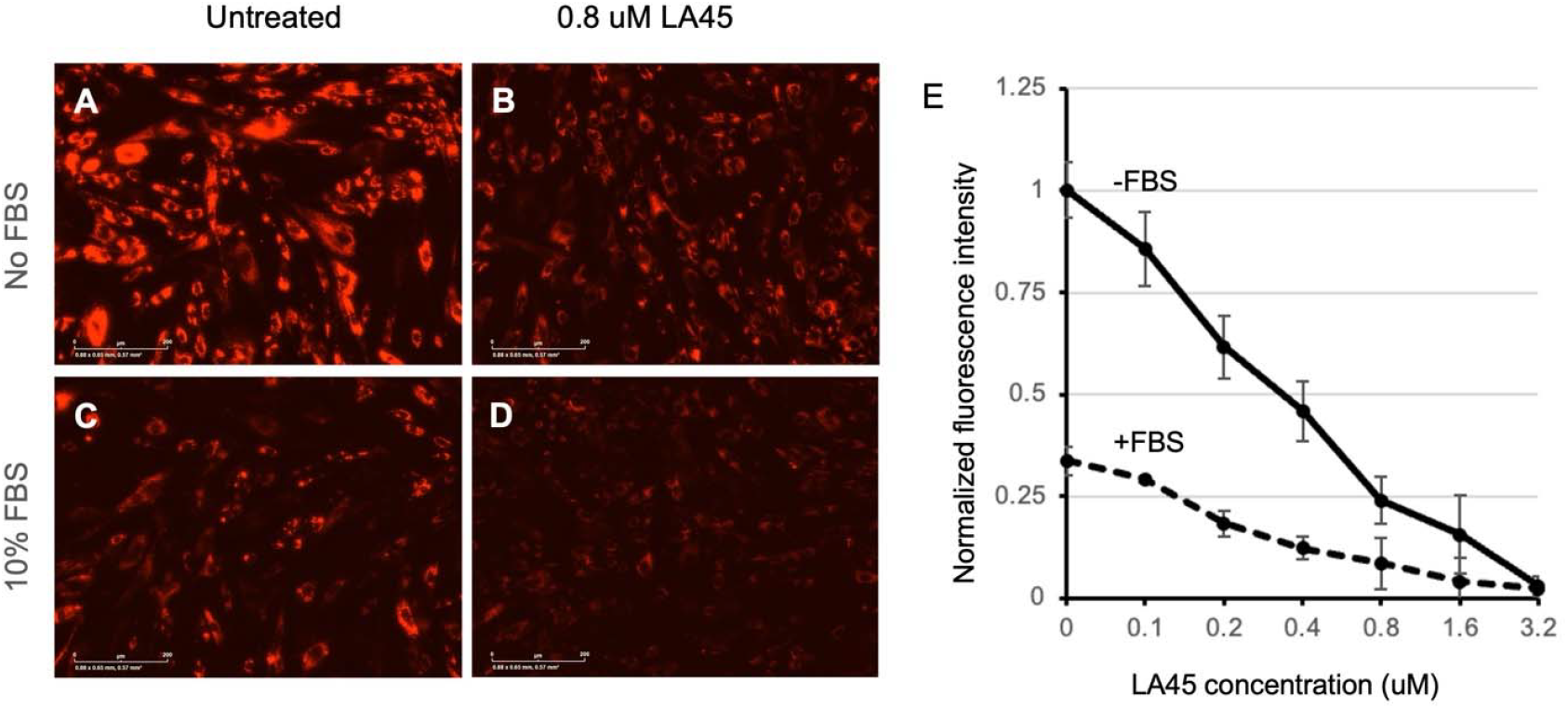
LA45 effectively inhibits ApoE4 endocytosis under physiological serum conditions. (A-D) Fluorescence microscopy of ApoE4 uptake: Representative images displaying the internalization of ApoE4 lipoparticles under serum-free (No FBS) and serum-containing (10% FBS) conditions. Panels A and C represent the untreated controls. Panels B and D depict cells treated with 0.8 µM LA45, showing a marked visual reduction in fluorescent lipoparticle accumulation compared to their respective untreated controls. (E) Quantitative dose-response curve: The graph plots the normalized fluorescence intensity against increasing LA45 concentrations (ranging from 0 to 3.2 µM). The solid line illustrates the inhibition trajectory in the absence of serum (-FBS), showing a steady, dose-dependent decline from the maximum baseline. The dashed line represents the 10% FBS condition (+FBS), revealing that although the initial baseline of ApoE4 uptake is intrinsically lower when serum is present, the LA45 peptide maintains its antagonistic functionality, effectively suppressing endocytosis down to negligible levels at 3.2 µM.

Representative fluorescence microscopy images clearly illustrate the peptide’s impact under both culture conditions. Under serum-free conditions (-FBS), untreated cells exhibited massive intracellular accumulation of ApoE4 lipoparticles (Figure 6A). Treatment with 0.8 µM LA45 resulted in a marked visual reduction in this lipoparticle accumulation (Figure 6B). When cells were cultured in media supplemented with 10% FBS (+FBS), the initial baseline of ApoE4 uptake in untreated cells was intrinsically lower compared to the serum-free baseline, likely due to native competition from serum lipoproteins for receptor binding (Figure 6C). Nevertheless, the addition of 0.8 µM LA45 still visibly and effectively suppressed the remaining endocytic uptake (Figure 6D).

Quantitative analysis of the dose-escalation matrix further validated these visual observations (Figure 6E). In the serum-free control, LA45 robustly inhibited ApoE4 endocytosis, demonstrating a steady, dose-dependent decline in normalized fluorescence intensity from 0 to 3.2 µM, achieving near-complete inhibition at the maximum dose. In the 10% FBS condition, the starting normalized fluorescence intensity was reduced to approximately 35% of the serum-free maximum. Despite this lower baseline and the highly competitive serum environment, the LA45 peptide maintained its potent antagonistic functionality. It effectively suppressed the remaining endocytosis in a dose-dependent manner, driving the intracellular accumulation down to negligible levels at the 3.2 µM concentration. This confirms that the LA45 minireceptor retains robust target engagement and blocking efficacy even under physiological constraints.

## 4. Discussion

The complex and pleiotropic role of ApoE4 in Alzheimer’s disease presents ongoing challenges for developing targeted therapeutics. While significant clinical efforts have focused on addressing downstream pathological hallmarks such as amyloid-beta plaques and neurofibrillary tangle, our research explores a potentially critical upstream cellular pathway: the LDLR-mediated endocytosis of ApoE4. The structure of ApoE4, specifically its unique intramolecular domain interaction between the N-terminal and C-terminal regions (Arg61 to Glu255) restricts its conformational flexibility (Dong et al., 1994). This structural tethering may favor a more compact conformation that affects its lipid-binding capacity. Consistent with our FPLC/SEC results (Figure 1C), which demonstrate an isoform-dependent shift in the size of ApoE lipoparticles, previous studies have shown that ApoE4 tends to form smaller, less lipidated particles compared to the larger complexes typically formed by ApoE2 and ApoE3 (Gong et al., 2002; Hatters et al., 2006). Despite this reduced lipidation, the compact orientation of ApoE4 may stabilize the receptor-binding region, contributing to a higher binding affinity for LDLR compared to ApoE3 and ApoE4 (Zhong & Weisgraber, 2009; Yamamoto et al., 2008; Guo et al., 2025). These smaller, less lipidated particles, combined with a strong LDLR binding affinity, facilitate ApoE4 acting as an efficient vehicle for pathogenic endocytic overload.

The strong isoform-dependent endocytic profiles observed in our primary CNS models are generally consistent with recent findings in immortalized cell lines. A study by Guo et al. (2025) using HAP1 and H4 cells suggested that LDLR, rather than LRP1, primarily drives ApoE3 and ApoE4 uptake over a 24-hour period. Their data indicated that while ApoE3 and ApoE4 lipoparticles are internalized and trap LDLR in endolysosomal compartments, ApoE2 lipoparticles largely evades this uptake, thereby preserving surface receptor levels. Translating these receptor-ligand dynamics into primary astrocytes, iAstrocytes, and human microglia, our tracking data support the idea that an LDLR-driven isoform bias contributes to the heightened intracellular accumulation of ApoE4 lipoparticles across these relevant CNS cell types, whereas ApoE2 entry remains comparatively minimal (Figure 4).

The continuous influx of lipid cargoes into the endo-lysosomal system could serve as a contributing factor to cellular stress. Enlarged endosomes are considered one of the earlier neuropathological observations in Alzheimer’s disease, frequently noted prior to significant amyloid deposition (Cataldo et al., 2000; Nuriel et al., 2017). Our 24-hour live-cell assays model this process, illustrating how increased ApoE4-mediated lipid payloads might burden the lysosomal degradation machinery. This intracellular accumulation could potentially lead to lysosomal stress, the buildup of cholesteryl esters and lipid droplets in glia, lipofuscin accumulation, and the subsequent exacerbation of aggregated tau (Chan et al., 2012; van der Kant et al., 2019; Guo et al., 2025).

To address this potential endo-lysosomal burden, we explored the use of recombinant LDLR minireceptors as extracellular decoys. By intercepting ApoE4 before it engages native cell-surface receptors, we aimed to decrease ApoE-receptor interactions and alleviate stress on the endo-lysosomal system. A notable observation in our study is the target selectivity of these engineered minireceptors. The LA3456 construct demonstrated a capacity to bind ApoE4 lipoparticles while exhibiting minimal cross-reactivity with human serum LDL (Figure 2). Because systemic LDLR interactions regulate peripheral ApoB-mediated cholesterol transport (Goldstein & Brown, 2009; Wadhera et al., 2016), avoiding cross-reactivity with ApoB and human serum albumin is an important translational consideration. This selectivity provides a biochemical rationale for further investigating these constructs as leads that may minimize interference with peripheral ApoB-mediated cholesterol transport. The structural requirements for this targeted interception are reflected in our observed endocytosis inhibition hierarchy: LA3456 _≈_ LA345 > LA456 _≈_ LA45 >> LA34 (Figure 5). This functional ranking generally aligns with previous crystallographic and mutagenesis studies of the native LDLR, which indicated that ligand-binding repeats 4 and 5 (LA4 and LA5) form a core interface for ApoE recognition (Fass et al., 1997; Fisher et al., 2004). The relative ineffectiveness of the LA34 construct suggests that the complete LA4-LA5 tandem is likely necessary for functional ApoE4 antagonism. Additionally, the inclusion of flanking modules (such as LA3) in the longer constructs might confer enhanced conformational stability or cooperative binding avidity, which could help explain their lower estimated IC_50_ values (Figure 5).

Beyond LA module composition, we also investigated how molecular tagging location affects the activity of the minireceptors. In our dose-response assays, N-terminal tagging (such as the H6-SUMO-linker-LA45 and H6-linker-LA45 constructs) severely reduced the efficacy of the peptide. Even at extreme concentrations of 6.4 µM, the N-terminally tagged peptide failed to fully suppress ApoE4 entry. In contrast, when structural tags were fused to the C-terminus, creating the LA45-HA and LA45-V5 constructs, the inhibitory function was fully preserved. Quantitative assays across both primary and ApoE KO iAstrocytes confirmed that these C-terminally tagged minireceptors suppressed ApoE4 uptake just as efficiently as the unmodified LA45 peptide. This spatial constraint suggests that a free N-terminus may be important for maintaining the proper orientation or optimal conformational presentation of the LA4-LA5 binding interface.

The development of peptide therapeutics requires considerations of proteolytic stability in biological fluids. The sustained inhibitory activity of the LA minireceptors observed throughout our live-cell assays indicate a reasonable degree of structural stability. Our LC/MS-TOF and SDS-PAGE analyses confirmed that these recombinant peptides preserve their native architecture, characterized by three dense intra-module disulfide bonds per LA domain (Bieri et al., 1995). This rigid intra-module disulfide bonds may help protect the binding pockets, allowing LA45 to retain its antagonistic functionality even in the more complex, competitive environment of 10% FBS (Figure 6). Building upon this baseline of structural resilience, the LA345 construct appears to offer an optimal balance of properties. By combining the enhanced inhibitory activity of a three-domain architecture with sustained stability, LA345 emerges as a particularly promising candidate for future therapeutic development. While our findings establish the biochemical efficacy of the LA345 construct, the translational application of such peptide-based decoys will require sophisticated delivery strategies to navigate the blood-brain barrier and ensure sufficient CNS bioavailability.

In conclusion, our findings support the premise that ApoE4 toxicity is closely linked to a receptor-mediated endo-lysosomal burden. Targeted intervention using truncated, structurally optimized LDLR-LA minireceptors, particularly the stable and active LA345 construct, may offer a mechanism to selectively protect the cellular clearance machinery from ApoE4-induced stress. By intercepting this pathway before intracellular accumulation occurs, these minireceptors highlight a viable upstream target for therapeutic intervention in Alzheimer’s disease that merits further exploration.

## Supplemental

**Figure S1.**
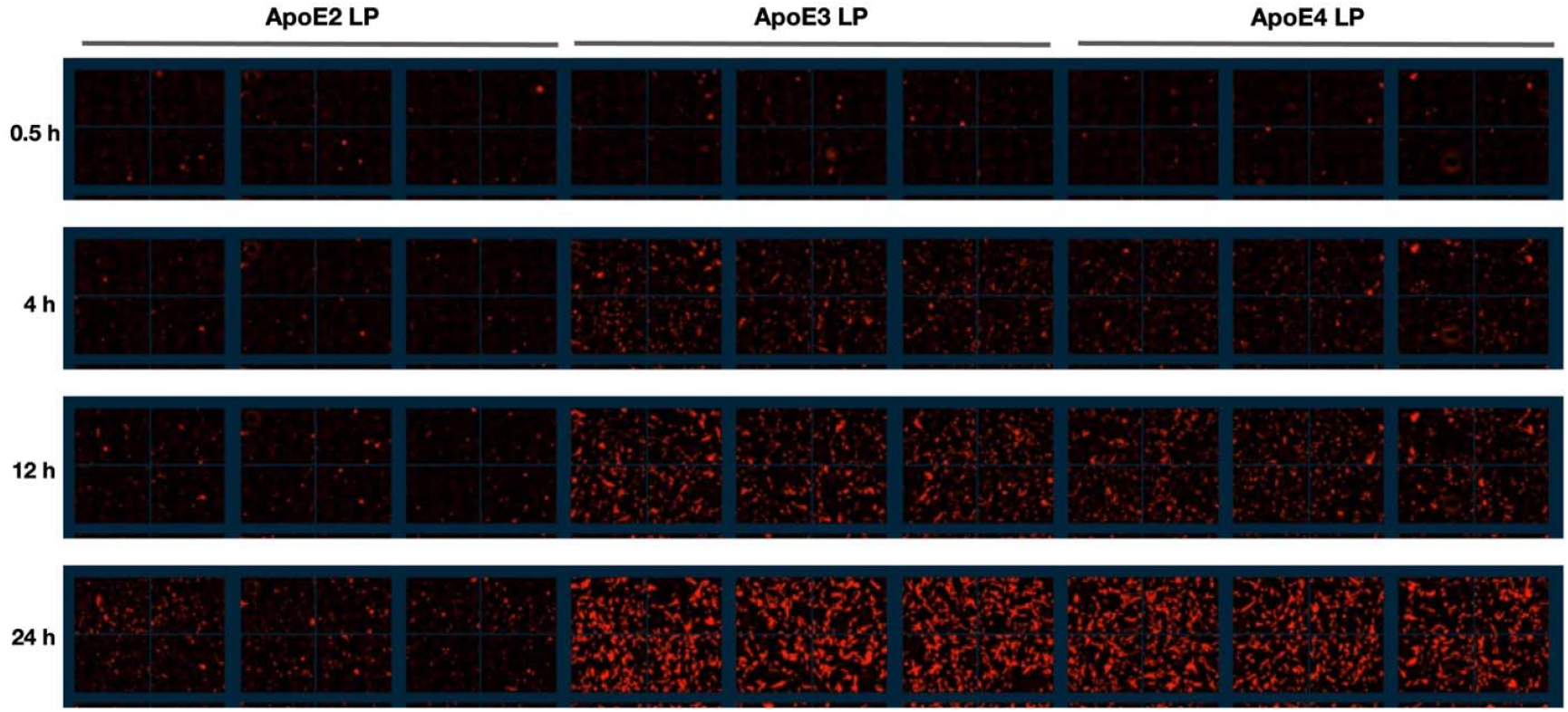
Detailed time-course and biological replicates of isoform-dependent ApoE lipoparticle endocytosis in primary human astrocytes. Representative live-cell fluorescence microscopy images tracking the internalization of 100 nM AF594-labeled ApoE2, ApoE3, and ApoE4 lipoparticles over a continuous 24-hour period. The assay was performed under standard serum-free conditions, matching the experimental condition of Figure 4, to establish baseline receptor-mediated uptake kinetics. Three independent replicate wells are displayed side-by-side for each ApoE isoform condition to illustrate the high reproducibility of the endocytic profiles. The chronological imaging matrix captures early endocytic events starting at 0.5 and 4 hours, progressing to massive intracellular accumulation at 12 and 24 hours for both the ApoE3 and ApoE4 variants. In contrast, the ApoE2 lipoparticles exhibit minimally detectable cellular uptake across all replicates throughout the entire duration of the assay. These chronological visual data directly corroborate the severe isoform uptake bias (ApoE4 _≈_ ApoE3 >> ApoE2) quantified in the main text.

**Figure S2.**
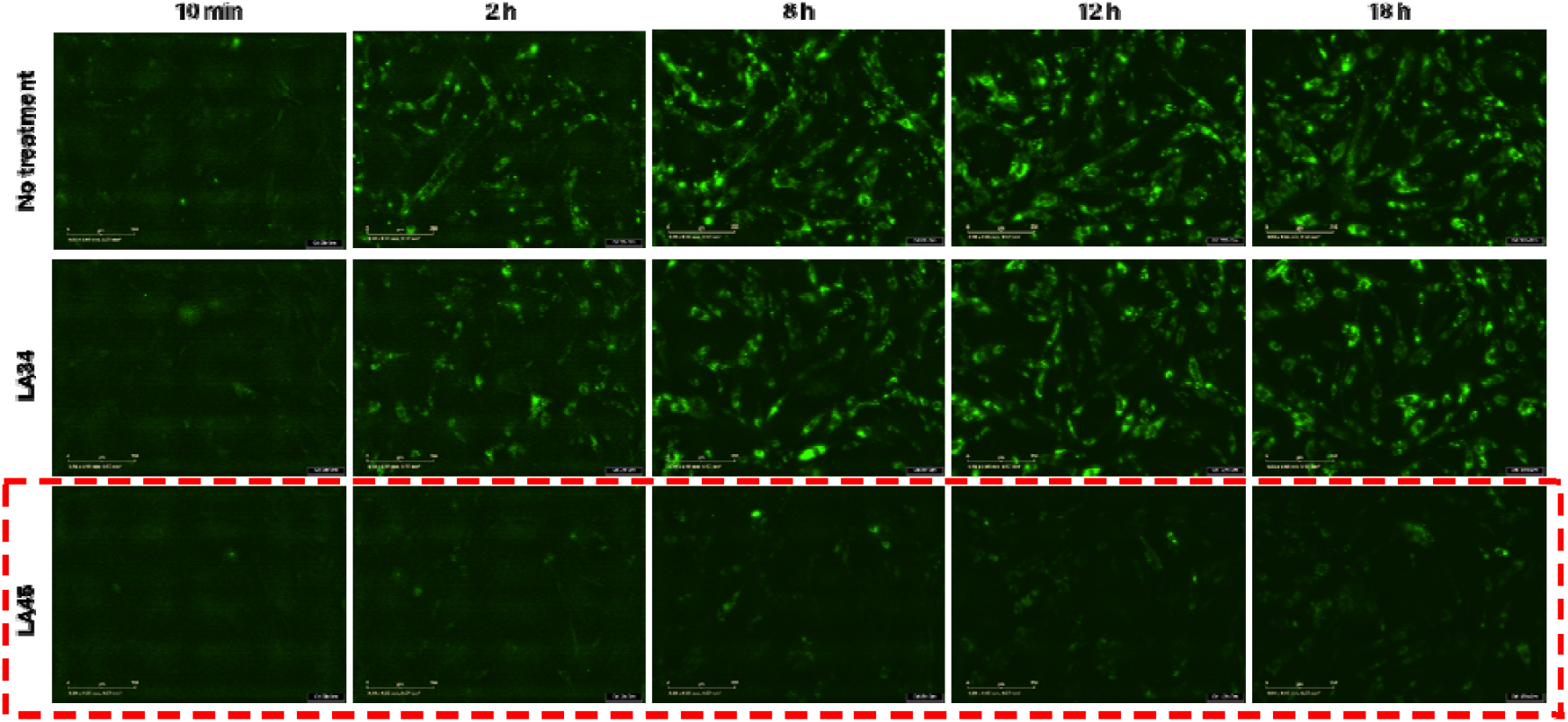
Time-course analysis of ApoE4 lipoparticle endocytosis and differential competitive inhibition by LA minireceptors. Representative live-cell fluorescence microscopy images tracking the internalization of 100 nM AF488-labeled ApoE4 lipoparticles over a continuous 24-hour period. The top row illustrates the robust baseline endocytic uptake and progressive intracellular accumulation of the ApoE4 lipoparticles in the absence of an inhibitor. The middle row depicts cells co-incubated with 1 µM of the LA34 minireceptor, demonstrating its ineffectiveness at inhibiting endocytosis, as the uptake profile remains comparable to the baseline. In contrast, the bottom row (highlighted by the dashed red box) demonstrates co-incubation with 1 µM of the LA45 minireceptor. The introduction of the LA45 decoy effectively intercepts the ApoE4 lipoparticles in the extracellular space, outcompeting native cellular receptors and resulting in a sustained reduction in endocytic volume across the entire time course. These visual data corroborate the quantitative competitive inhibition profiles detailed in the main text, highlighting the specific modular requirements for effective ApoE4 interception.

**Figure S3.**
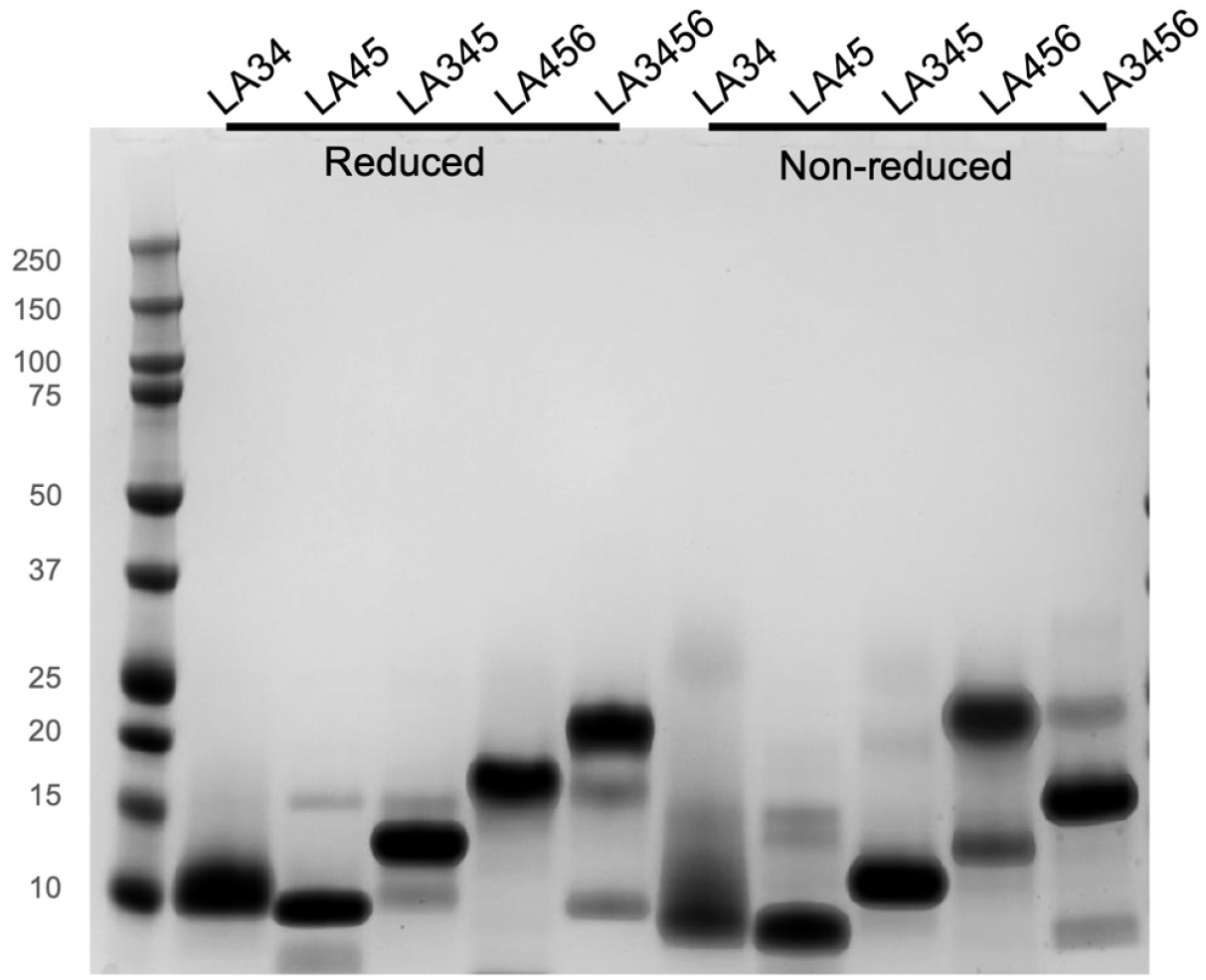
SDS-PAGE analysis of LA peptide variant stability. Recombinant LA peptides (LA34, LA45, LA345, LA456, and LA3456) were stored at 4°C for two weeks following size-exclusion chromatography purification to evaluate their structural stability. The gel illustrates the electrophoretic mobility of the samples prepared under reducing conditions (left five lanes, treated with 5 mM TCEP prior to loading) alongside the corresponding samples maintained under non-reducing conditions (right five lanes). The differential migration between the reduced and non-reduced states demonstrates that intact intra-module disulfide bonds maintain the non-reduced peptides in more compact conformations. Additionally, LA456 exhibited significant oligomerization relative to the other variants.

